# Lateralised memory networks explain the use of higher-order visual features in navigating insects

**DOI:** 10.1101/2024.11.26.625348

**Authors:** Giulio Filippi, James Knight, Andrew Philippides, Paul Graham

## Abstract

Many insects use memories of their visual environment to adaptively drive spatial behaviours. In ants, visual memories are fundamental for navigation, whereby foragers follow long visually guided routes to foraging sites and return to the location of their nest. Whilst we understand the basic visual pathway to the memory centres (Optic Lobes to Mushroom Bodies) involved in the storage of visual information, it is still largely unknown what type of representation of visual scenes underpins view-based navigation in ants. Several experimental studies have shown ants using “higher-order” visual information – that is features extracted across the whole extent of a visual scene – which raises the question as to where these features are computed. One such experimental study showed that ants can use the proportion of a shape experienced left of their visual centre to learn and recapitulate a route, a feature referred to as “fractional position of mass” (FPM). In this work, we use a simple model constrained by the known neuroanatomy and information processing properties of the Mushroom Bodies to explore whether the use of the FPM could be a resulting factor of the bilateral organisation of the insect brain, all the whilst assuming a “retinotopic” view representation. We demonstrate that such bilaterally organised memory models can implicitly encode the FPM learned during training. We find that balancing the “quality” of the memory match across left and right hemispheres allows a trained model to retrieve the FPM defined direction, even when the model is tested with other shapes, as demonstrated by ants. The result is shown to be largely independent of model parameter values, therefore suggesting that some aspects of higher-order processing of a visual scene may be emergent from the structure of the neural circuits, rather than computed in discrete processing modules.

**Author summary:** Many insects are excellent visual navigators, often relying on visual memories to follow long foraging routes and return safely to their nest location. We have a good understanding of the neural substrates supporting the storage of visual memories in ants. However, it is still largely unknown what type of representation of visual scenes underpins the functions of visual navigation. Experimental studies have shown ants using “higher-order” features as part of navigation, that is features that are extracted across the whole extent of a visual scene. Using an anatomically constrained model of the insect memory centers, we address the question of whether the use of higher-order visual features may be emergent from the overall architecture of the vision-to-memory pathways. We find that balancing the quality of left and right visual memory matches provides an explanation for some higher-order visual processing and visual cognition shown in experiments with ants. Overall, this constitutes a contribution to our understanding of visual cognition and the processing of visual scenes used in navigational tasks. We additionally postulate a novel mechanism ants may use to navigate, which is supported by the bilateral structure of the insect brain.

## Introduction

Insects often rely on visual memories to navigate routes within their environment [1–4] and also to relocate goal locations [5–8]. Models of view-based homing and navigation have shown that “snapshot matching” – the comparison of the current view with a single or multiple views committed to memory – can be used for homing [6, 9–11] or route following [12–15]. Whilst we have a good understanding of the brain regions supporting visual memories in ants [16–18], much less is known about the visual processing that precedes the memorization of visual scenes [19]. It is also largely unknown how the bilaterally organised structure of the visual and memory circuits contributes to the functions of visual navigation [20, 21]. In this work, we consider behavioural experiments that explored several aspects of visual scene perception and visual cognition in ants [22], and explore how the known bilateral neuroanatomy of the visual and memory circuits can help in explaining these experimental results.

Two overarching theories of image processing are the holistic and feature-based approaches, which respectively favour a mostly retinotopic image representation and a parameterized image representation [23]. Whilst much behavioural evidence is consistent with insects learning and using almost “raw” retinotopic images for navigation [19], several experimental studies suggest that insects extract higher-order parameters derived from the whole visual scene. For example, it has been shown that insects can use the centre of mass (CoM) of large shapes [24–27]. Theoretical work has proposed that ants could extract rotationally invariant features based on Zernike moments [28, 29]. In this work, we focus on the experimental findings of Lent et al (2013a), which show that ants can learn a heading direction using the proportion of a shape that lies to the left of their visual center, a proportion coined ‘fractional position of mass’ or FPM for short (Fig 1A).

**Fig 1.**
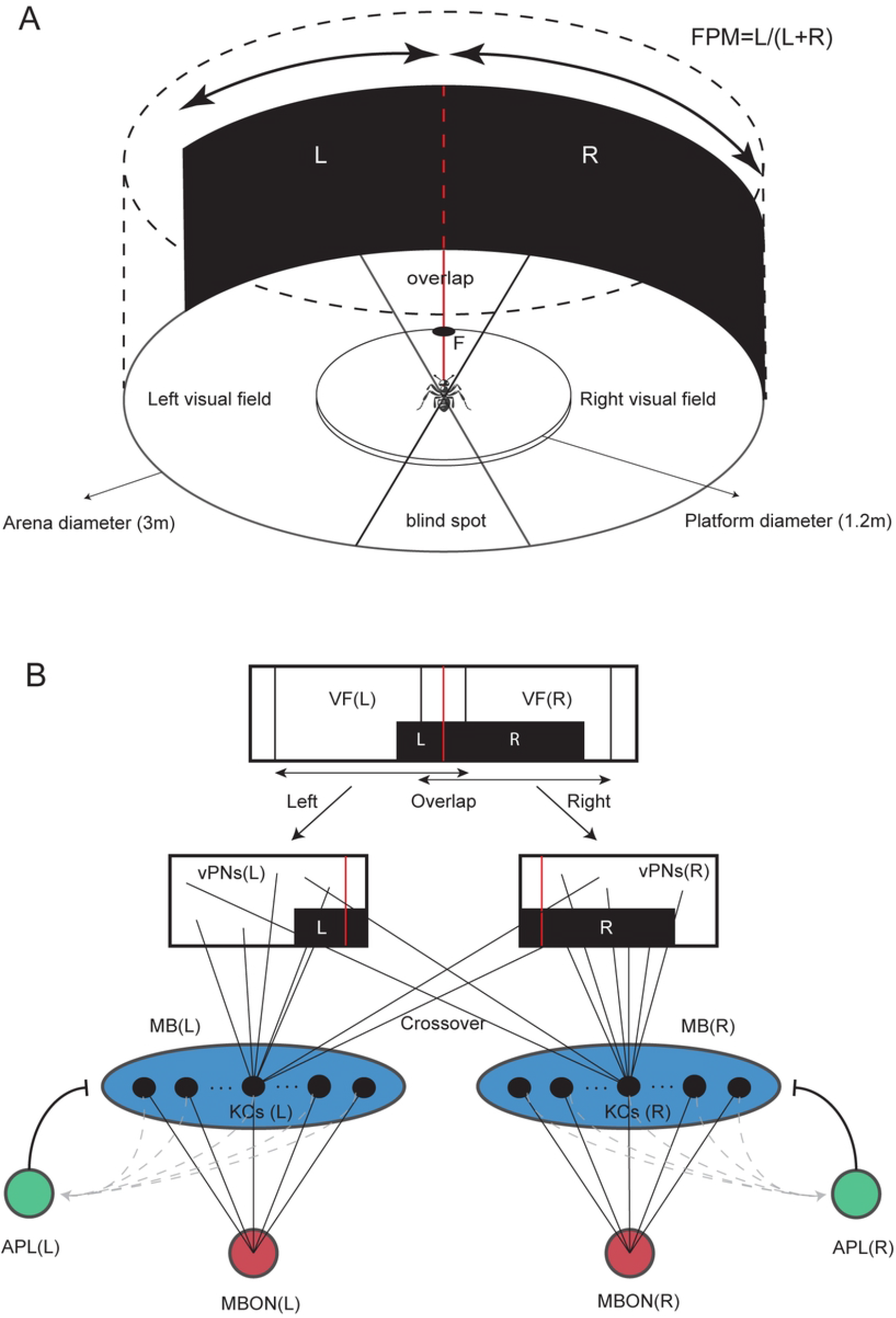
Experimental setting and network architecture. (A) Experimental Setting. The experimental setting of Lent et al (2013a) consists of a cylindrical arena with 3m diameter and 1.8m high walls. The feeder location is denoted with an F in schematic. The proportion *L/*(*L* + *R*) of a shape occurring on the left of the direct path to the food (center red line) is referred to as the fractional position of mass (FPM). In the diagram, we show an ant at the center of the arena (not to scale), as well as it’s left and right visual fields. In modelling, we assume left and right visual fields have a range of 180 degrees with some of their field of view overlapping in the front (overlap region), leading to a blind spot at the back. (B) Network Architecture. We model the ant’s memory circuit using an artificial neural network (ANN) with anatomical and information processing features drawn from properties of the insect Mushroom Bodies (MBs). The visual scene in panoramic form is split into left and right visual fields (VFs) by cropping at the appropriate azimuthal angles. The visual projection neurons (vPNs) are taken to be the pixel values of the visual fields. Each of the left and right Mushroom Body compartments contains *N*_*KC*_ Kenyon Cells (KCs). vPNs are connected to KCs randomly so that on average KCs have *K* connections (connections shown only for one KC). The left and right MBs are each recurrently connected to an Anterior Paired Lateral neuron (APLs) which serves to normalise MB activity. Each of the MBs is also fully connected to a Mushroom Body output neuron (MBONs).

More specifically, Lent et al (2013a) trained ants to head to a feeder within a cylindrical arena, placed at given location relative to a large shape (visual context). The ants were subsequently tested with different shapes, to see how an imperfect memory match would translate into behavioural decisions. The results of the experiments showed that ants tend to keep the same proportion of a test shape left of their visual center as they had experienced in training. In other words, ants navigate to the same FPM in test scenarios as the one they are trained with. These findings suggest that ants can extract higher-order features computed over the whole extent of a visual scene to learn and recapitulate routes, in this case the FPM of a large shape. When ants were trained with composite shapes, the experimental results further suggested that ants segment the visual scene prior to computing the FPM, another example of higher-order visual processing and visual cognition.

Recent lesioning experiments with ants have shown that visual memories are supported by the Mushroom Bodies [17, 18], a highly conserved pair of structures in the insect brain known for their role in memory and associative learning [30, 31]. Therefore, in this paper, we explore whether the use of the FPM in navigating wood ants (and other, higher order visual cognitive processes such as shape segmentation) can be explained by a simple model leveraging the known anatomy and information processing properties of these memory centers. We assume a simple “pixel-wise” retinotopic image representation, and propose that, nevertheless, indicators of higher-order visual processing emerge from the functional properties of the downstream memory circuits. Building on recent modelling approaches to visual navigation considering the bilateral organisation of the insect brain [20, 21, 32–34], our proposed model is equipped with two ‘eyes’ and two Mushroom Bodies (MBs) each with it’s own output neuron. We allow visual information from the left and right eyes to project into both the left and right MBs, in line with neuroanatomy [35–37]. However, we introduce a stochastic bias in favour of ipsilateral visual projections, so that each MB specializes in processing information from its ipsilateral field of view (lateralisation). We show that, with sufficient bias, a signal for the FPM defined direction is implicitly encoded within the model, and that this result is largely independent of parameter values. In so doing, we demonstrate that some higher-order processing of the visual scene may be emergent from the bilateral neuroanatomy of the circuits, rather than computed in discrete processing modules.

## Results

### A simple bilateral memory model captures FPM results

In this work, we wanted to explore whether the use of the FPM in navigating ants [22] could result from the bilateral structure of the visual and memory circuits in the insect brain. To do this, we consider a simple agent with an ant inspired visual navigation circuit, within a simulated version of the cylindrical arena from the Lent et al., (2013) experiments (Fig 1A). The memory model is implemented as an artificial neural network, retaining the key anatomical and functional properties of the insect Mushroom Bodies (Fig 1B). The visual scene is split into left and right hemispheres prior to projection into the (left and right) Mushroom Bodies (MBs), with a bias favouring ipsilateral connections. Implementational details for each neuron type and the network learning rule are described in detail in the Methods section, and are similar to other approaches taken in the field (e.g., [38]). For a given training shape, the model is trained on a direct linear path from the center of the arena to the feeder location, with views facing forward. The trained model is subsequently tested with different shapes, and the informational properties are studied using the outputs of the independent left and right MBONs (which constitute novelty signals) taken across all possible rotations of the agent. An example of left and right rotational novelty signals, alongside their sum and difference are given in (S1 Fig).

In Fig 2, we show the results for the first set of shapes, which consist of large rectangles and trapezoids on white background. In Lent et al (2013a) the distributions of aiming points of the ants are calculated as the distributions of the end points of saccade-like-turns (SLTs) as projected onto the arena wall (row (iii)). SLTs are periods of very high rotational velocity, shown in [39] to be visually guided turns correcting the facing angle towards a goal direction. Therefore, to model the heading distributions, we needed a way to generate a distribution of simulated saccade end points from the outputs of our computational memory model. Whilst it has been shown that the heading directions of ant navigation can often be modelled by minimizing a visual novelty signal [12, 14, 15], our additional modelling insight is that ants, given bilaterally organized memories, might also need to balance left and right novelties. Our approach therefore assumes that both the proximity of the novelty sum to it’s minimum value (low novelty signal), and the proximity of the novelty difference to 0 (lateral balance signal) have equal importance in generating goal directions. To ensure that the observed results are a robust property of our model architecture and not the result of parameter tuning, we display the simulated distributions pooled across a broad range of parameter combinations (see Methods).

**Fig 2.**
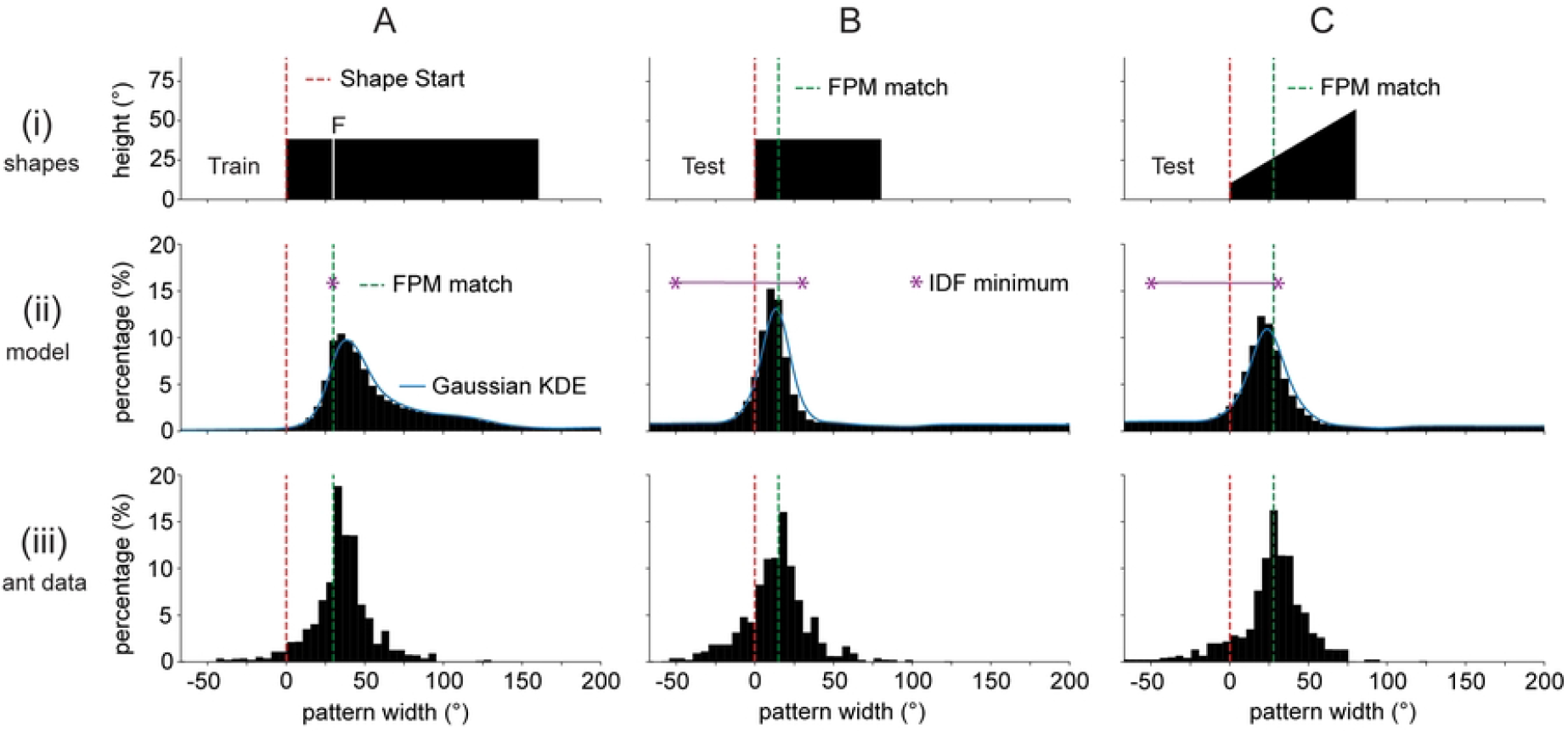
Results for basic FPM demonstration. Column (A). (i) The training shape for the first set of experiments consists of a rectangle of width 160° and height 38° (stimulus angles measured from center of arena). The white line represents the feeder location. (ii) Distribution of model derived headings for the training shape tested against itself. The red dotted line and green dotted line represents the start of the shape and the FPM-match direction respectively. The blue curve is a KDE approximation to the simulated distribution, with gaussian kernel and bandwidth of 5°. The purple asterisk(s) denotes the minimum of the Image Difference Function (IDF), or the angular range over which the IDF is flat. (iii) Experimental goal direction distributions inferred from Lent et al (2013a). Goal directions are measured using the facing direction of saccade-like body turns (SLTs) as projected onto the arena wall [22, 39]. (B). Test shape (i), model distribution (ii) and original ant data (iii) for a test shape of a 80° by 38° rectangle. (C). As for (B) but for a trapezoid of width 80°, with heights of 10° and 57° on the left and right sides respectively.

In Fig 2 row (ii), we show the simulated distributions for the first series of experiments in Lent et al., (2013) alongside the experimentally obtained distributions in row (iii). As can be observed in the plots, the simulated distributions capture the major mode of the experimental findings across all three examples. We use the distance between simulated modes and true modes inferred from Lent et al (2013a) as a measure of performance. Across the three test shapes, the distance between modes between the simulated and experimental distributions is below 6.1°. If we evaluate the distributions for all parameter combinations and shapes separately, we get an average distance between modes of 5.0° ± 3.5° (mean ± std). The average performance of the model for each of the parameter settings taken independently are shown in (S2 Fig Panel A) which demonstrate that the model performs well across all tested parameters. Overall, these results show that the simulated distributions capture the experimentally obtained distributions for simple shapes, and that the results are robust against changes in parameter settings. This suggests that the occurrence of a major mode at the FPM-match direction for this class of shapes is a general principle resulting from the bilateral architecture of the model rather than a specific tuning of its parameter values.

The success of the model in capturing the pattern of results from the biological data is in large part due to the balancing of left and right novelties from the independent MBs. Indeed, a careful analysis shows that there is no signal for the FPM-match direction within the left and right rotational novelty curves taken alone (S1 FigA-B), nor is there a consistent signal for the FPM-match direction within the rotational novelty sum (S1 FigC). This finding mirrors the original finding of Lent et al (2013a) who reported that Image Difference Function (IDF) models are inadequate to describe their experimental outcomes. This is because for test shapes smaller than the train shape, the IDF flattens over a large angular interval (in Fig 2, the purple line represents region where IDF is minimal), which leaves the goal direction under-determined. In our present framework, we can think of the novelty sum as a kind of IDF, which measures the overall match of the train and test shapes, which therefore does not consistently contain a signal for the FPM defined direction. On the other hand, equalizing left and right novelties resolves the indeterminacy, and helps in predicting the location of the experimental modes (S1 FigD). In the next section, we study the model in a theoretical setting, and provide an explanation for why we expect left and right novelties to equate at the FPM-match direction for some train and test shape combinations.

### FPM-match direction balances left and right novelties in theoretical analysis for nested shapes

In this section, we provide a theoretical explanation as to why we expect left and right novelties to equate at the FPM-match direction for a variety of train and test shape combinations. To do so we use the bilateral memory model (as described in Methods), with the simplifying assumption that there is a pure lateral split (no crossover connections, and no visual overlap). We further assume that the test shape can be contained within the train shape (nested) so that we can separate the shape overlap regions into 4 areas: train-left(X), test-left(Y), train-right(W), test-right(Z) (Fig 3A). The first step is to notice that when both train and test images are aligned at the same FPM value, the following equation holds:

**Fig 3.**
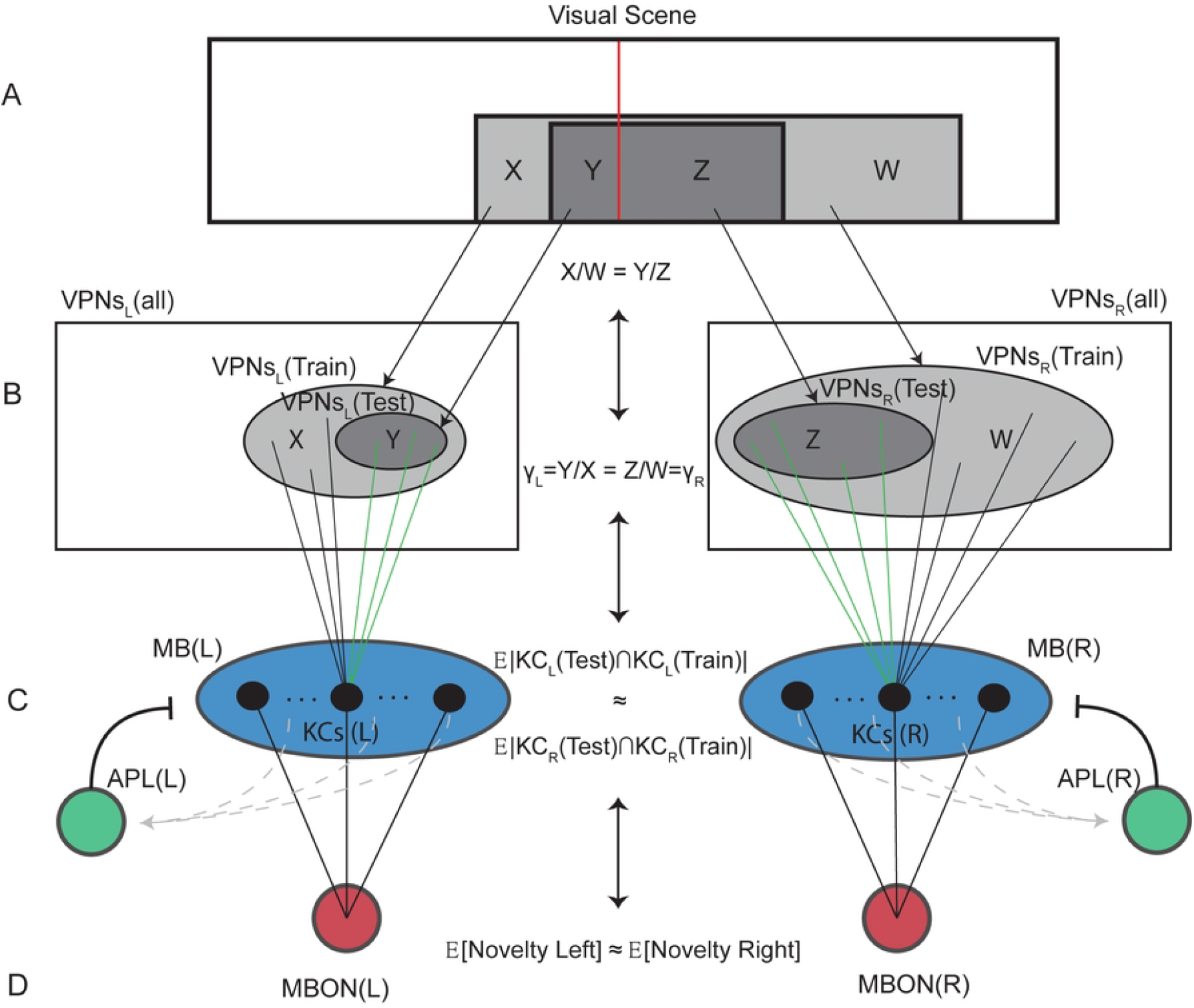
The theoretical basis for the encoding of FPM by the bilateral memory model. This figure represents the signal flow as the information progresses through the network layers. (A) Train and test shapes (schematic) are shown in light and dark grey respectively. The panoramic visual scene is split into left and right visual fields (VFs) by a center line (red line). This splits the two rectangles into two parts which we denote as X and W for the train shape and Y and Z for the test shape. Since we assume the two rectangles are aligned at the same FPM, we have 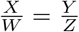. (B) The visual projection neurons split into left and right (vPNs). We show two embedded sets of vPNs for the neurons that would fire for train and test shapes respectively, drawn as ovals in light and dark grey. The equality of FPM values rearranges into an equation that equates laterally segregated informational quantities 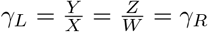. (C) The two Mushrooom Body networks (MBs), each with an anterior paired lateral neuron (APL). We show connections only for one Kenyon Cell (KC) and restrict our focus to those Kenyon Cells that fired for the training shape, with connections uniformly distributed on the training vPN sets. Connections that are contained (“captured”) in the test firing vPN set are shown in green. We also show the expected KC intersection sizes approximately equating across hemispheres. (D) Left and right MBONs which compute the novelty of the bilateral portions of the test shape, relative to the learnt shapes. We show the left and right novelties equating approximately. Equivalence lines running up and down the center of the image indicate that the chain of causality in the equations can be seen as running up the signal flow as well as down.

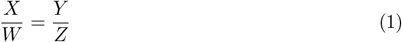

If we define *γ*_*L*_ and *γ*_*R*_ to be the proportion of the train shape covered by the test shape in the left and right visual fields respectively (Fig 3B). Then Equation 1 can be rearranged into:

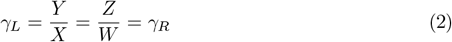

So far, we have shown an equivalence between the alignment of the images at the same FPM, and the test shape covering an equal proportion of the train shape in the left and right hemispheres. The second step is to justify why we expect left and right novelties to equate when *γ*_*L*_ = *γ*_*R*_.

To do this, we start with an expression for the expected (Left) test novelty, subject to the model having learnt the train image. Because of the learning rule used (see Methods), we can relate the novelty of the test shape to the expected number of Kenyon Cells that fire on both train and test images (Fig 3C).

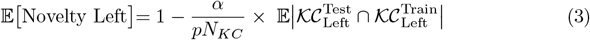

In the above expression, *α* is the learning rate used in training, *p* denotes the probability of a Kenyon Cell firing and *N*_*KC*_ is the number of Kenyon cells in one hemispheric MB. We use calligraphic notation 𝒦𝒞 to denote sets of Kenyon Cells rather than a single cell. By manipulating the above expression (using an identical distribution of KCs), we can rewrite it in terms of the conditional probability of a single Kenyon Cell firing on the test image, given prior knowledge that it fired on the train image.

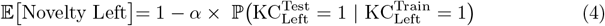

From the above formula, we deduce that left and right novelties equate when the conditional firing probabilities equate on the left and right sides. For a Kenyon Cell to fire, it needs to have an appropriate number of coincident inputs from firing vPNs, relative to the population average (e.g., exceed (1 − *p*)^*th*^ quantile for our APL model). Given the prior knowledge that the Kenyon Cell fired in response to the training image, the Kenyon Cell also fires in response to the test image if sufficient of the train connections also coincidentally connect within the test firing vPN set (displayed as green connections in Fig 3C). Because the pattern of connections is assumed to be random, the connections to vPNs that fired at train time are randomly distributed homogeneously on the train firing vPN set. As a consequence of this fact, each connection with a firing vPN is contained (“captured”) within the test firing vPN set, with a probability given by the covering proportion *γ*.

A low value of *γ* means almost no connections are captured and a high value of *γ* (close to 1) means almost all connections are captured. For the test firing vPN set to capture the same (expected) number of connections in both left and right hemispheres relative to the population averages, the left and right covering proportions must be equated. Capturing the same proportion of connections on both sides ensures the conditional distributions get the same “edge” over the population distributions, which is why we expect *γ*_*L*_ = *γ*_*R*_ to balance out left and right conditional firing probabilities, and therefore also image novelties (Fig 3D).

In light of the theoretical explanation as to why left and right novelties balance at the FPM-match direction, we do not expect the result to be sensitive to parameter values. Indeed, running the computational model for many simple shapes and parameter settings shows that in almost all cases the rotational novelty difference has a zero-crossing at the FPM-match direction (S3 Fig). This includes weight initialization decisions, for instance we can initialize all vPN to KC weights randomly, all KC to MBON weights randomly, or all network weights randomly, and the rotational novelty difference still has a zero-crossing at the FPM-match direction in all three cases (S3 Fig row (vi)). The parameters that are the most likely to shift the location of zero-crossings are crossover and overlap parameters. Crossover values close to 0.5 lead to unstable outputs, as the bilateral difference signal becomes weak before flattening at 0.5 (S3 Fig row (iv)). High overlap values can considerably shift the location of zero-crossings, especially for smaller test shapes (S3 Fig row (v)).

### Modelling assumptions also explain experimental results for more complex shapes

In the previous two sections, we analysed cases where test shapes could be mostly contained within the train shapes. Following the second set of experiments by Lent et al (2013a), we consider more complicated interactions between training and test shapes composed of triangles and trapezoids. Whilst the theoretical argument does not apply to these train and test shape combinations, because test shapes cannot be contained within the train shapes, we will show that the same methodology for simulating goal direction distributions can still provide a strong heuristic for the location of modes in the experimental heading data.

The train shape used in this set of experiments is a 160° wide scalene triangle (Fig 4A). The first test shape (B) consists of the training shape reflected horizontally and the second test shape (C) consists of a 160° wide trapezoid. As can be observed in Fig 4 row (ii), the simulated distributions capture the major mode locations of the experimental distributions displayed in row (iii). Across the three examples, the simulated distributions have a distance between modes below 8.7°. If we instead split the analysis over all parameter values and shapes, the average distance between modes is 7.3°± 5.7° (mean ± std). The average performance of the model for each of the parameter settings considered are shown in (S2 Fig Panel B). These results show that the model still robustly estimates experimental mode locations.

**Fig 4.**
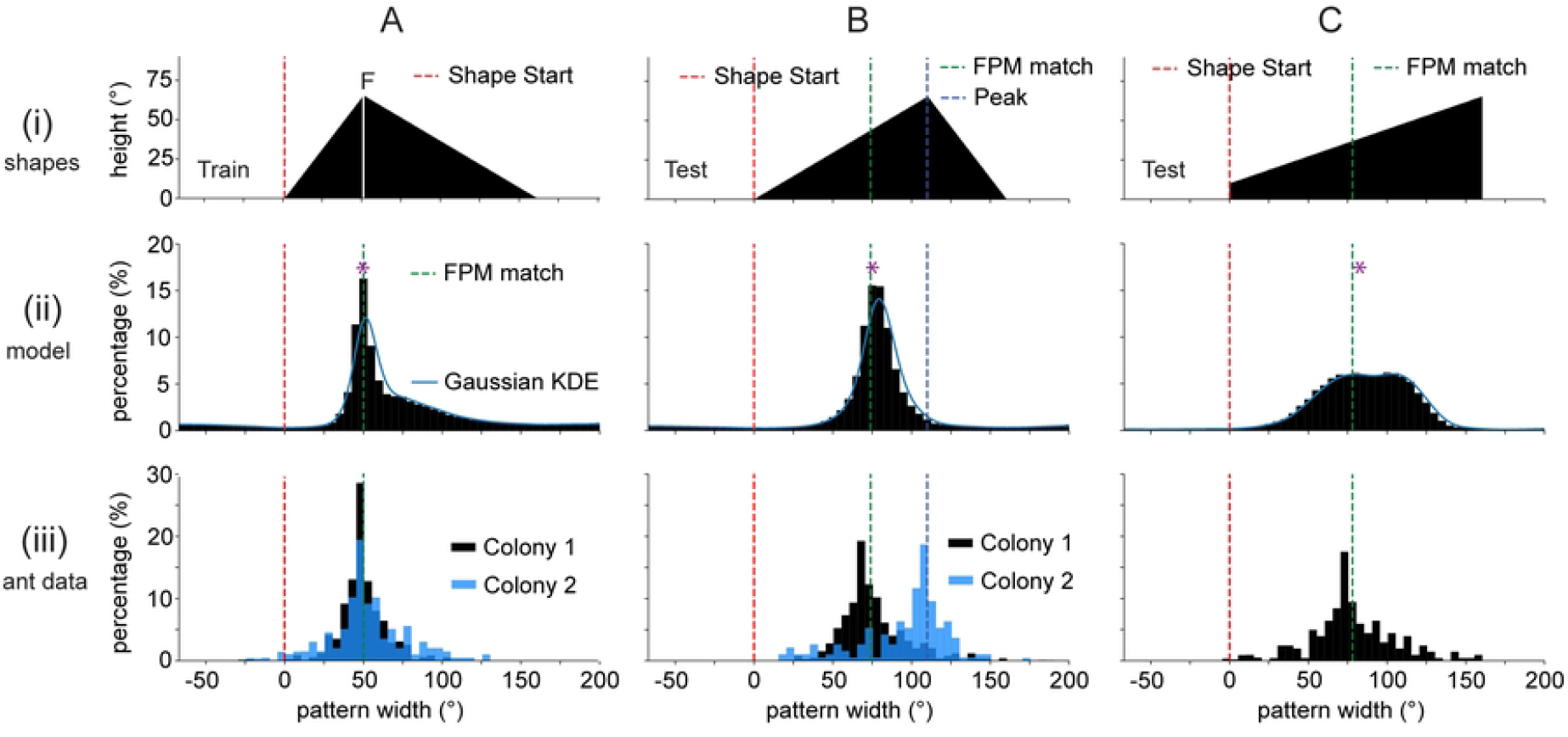
Results after training with a triangular shape. Column (A). The training shape for the second set of experiments consists of a scalene triangle with width 160° and height 65°, and summit 50° to the right of the shape starting point. Column (B). The first test image is the train image reflected horizontally. The blue dotted line represents the location of the peak. Column (C) The second test image is a trapezoid of width 160°, with heights of 10° and 65° on the left and right sides respectively. Row (i). Schematic depiction of the train and test images. Row (ii). Simulated saccade-end-point distributions. The purple asterisk denotes the minimum of the Image Difference Function (IDF). Row (iii) Experimental data inferred from Lent et al (2013a). For the train shape and first test shape, the data for two different colonies is displayed (Colony 1 and Colony 2) with colors black and blue respectively. Other Figure conventions as in Figure 3.

In this set of experiments, two different colonies were reported to have different behaviours on the second test shape, as displayed in the two diverging distributions (black and blue) of Fig 4B(iii). The distribution of heading directions for the second colony (Colony 2) is centered below the peak of the triangle instead of the FPM-match direction. Whilst the results clash with the outputs of our computational model displayed in Fig 4B(ii), these experimental results admit a natural interpretation within this modelling framework. If we imagine train and test shapes aligned on their peaks, then the two views are horizontal mirror reflections of each other around the center line. As a consequence of this, the comparison of the train and test shapes in the left and right hemispheres will result in equal matches, regardless of which method one chooses to compare the shapes. So, if the ants only memorize a single view from the center of the arena, the peak would define the angle where we expect left and right novelties to balance.

In practice, we do not expect ants to memorize a single view from the center of the arena, because ants are trained over a 0.6*m* path to the feeder location (Fig 1A). Given that the image deforms and grows as the ant moves closer to the shape, experimental results (in one case) and the computational model predict the FPM-match direction as the location for the major mode of saccade end points. However, if the memory of the path were biased towards the first few views (due to some unknown mechanism), then we would expect that left and right novelties equate at the peak-defined direction, providing a tentative explanation for the bifurcation in colony behaviors.

### Results for composite shapes, a tentative explanation of segmentation

Further experiments by Lent et al (2013a) consisted of training with a composite shape, made up of two abutting triangles (Fig 5). Across three sets of experiments (I, II and III in Fig 5) the same training shape is used with different feeder locations. Specifially, the feeder location shifts rightward with increasing numerals. The experimental results for this set of shapes suggest that ants perform visual shape segmentation, another higher-order processing of the visual scene, when the feeder direction is close to the centre of a component of the composite shape rather than the centre of the entire composite shape.

**Fig 5.**
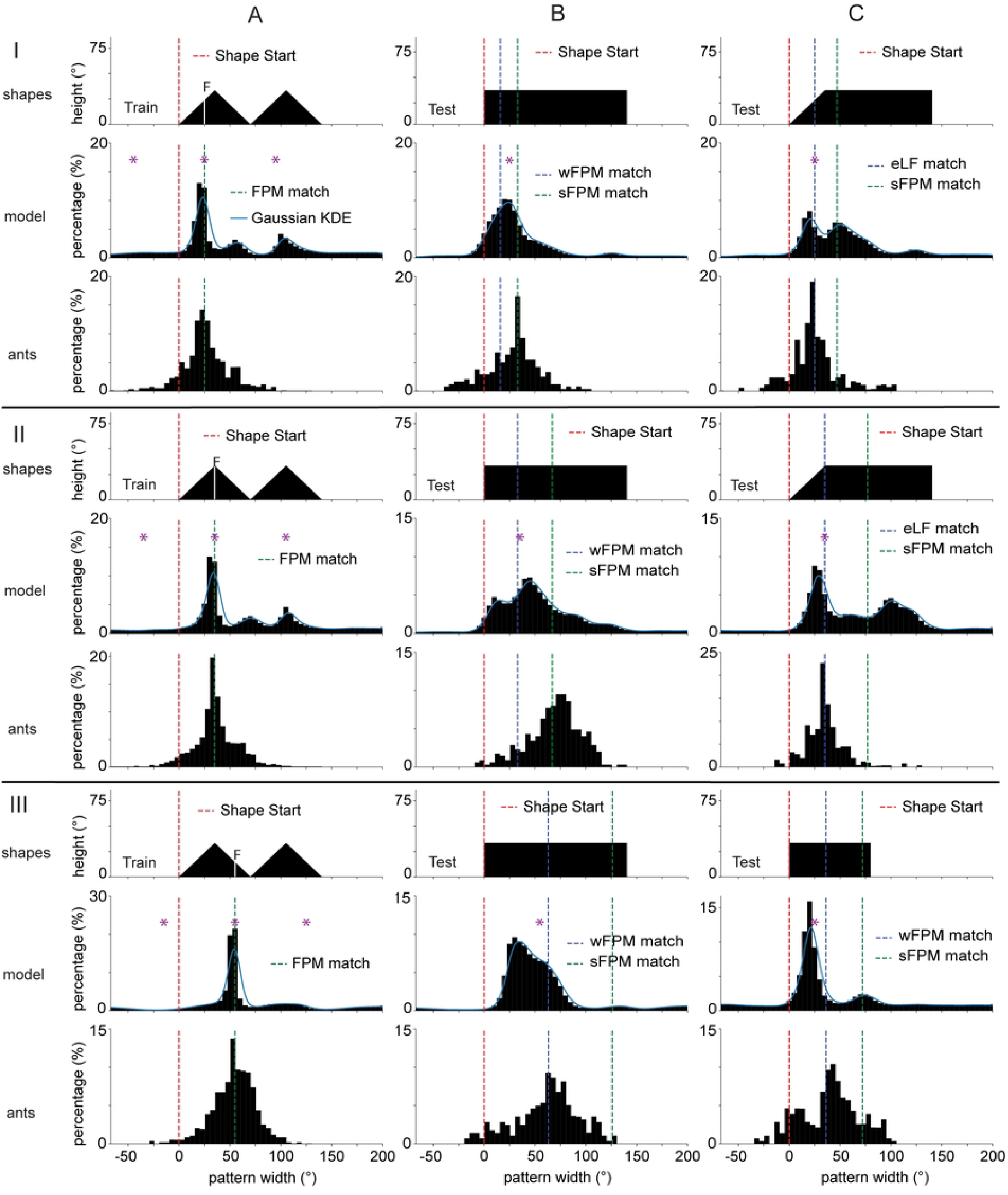
Results after training with composite shapes. This figure displays the results for three separate sets of experiments (I, II and III) where the training location is at different places within the same composite shape. Set I. The training shapes are a pair abutting triangles, each of which is isosceles with base 70° and height 35°. The feeder location is 25° rightwards of the shape start location. Set II. Same training shape as in previous set but the feeder is located at the center of the first triangle. Set III. The feeder is located 55° rightwards of the shape start location. Column (A). The results for the train shapes tested against themselves. Column (B). The test shape for all three sets is a rectangle 140° wide and 35° high. The blue dotted line represents the wFPM-match direction (FPM-match direction if we assume the FPM is computed across both triangles in training). The green dotted line represents the sFPM-match direction (FPM-match direction if we assume the FPM is computed across a segmented part of the image). Column (C). For sets I and II, the test shape is a trapezoid with width 140°, height 35°, and the top left vertex 35° rightwards of the start of shape. For these sets, we display a vertical dotted line at the eLF-match direction (direction where left slanted edge matches it’s retinal position between train and test runs). We also show a green dotted line at the sFPM-match direction, as described for previous set. For set III, the test shape is a 80° wide and 35° high rectangle. Across all examples, the purple asterisk(s) highlight the local minima of the rotational IDF. Other Figure conventions as in Figure 3.

Indeed, when ants are trained with a shape composed of two parts (e.g., two triangles), there are two possible ways that an FPM value might be extracted. The first is that the FPM is computed over the whole visual scene composed of two triangles, which is referred to as the whole FPM (wFPM). The second is that the FPM is computed over a segmented part of the scene (one triangle), which is referred to as the segmented FPM (sFPM). As observed in column (B) of Fig 5, when ants are trained to a feeder closer to the left of the composite shape, and tested with a wide rectangle, the goal direction distributions are centered around the sFPM (Fig 5 IB and IIB). This is indicative that the ants learned about heading directions relative to the leftmost triangle, and therefore suggests a shape segmentation mechanism (or attention-like process, see Discussion). On the other hand, if ants are trained to a feeder close to the center of the composite shape, the goal direction distribution, with the test shapes, are centered at the wFPM (Fig 5 IIIB), which is indicative that in this case no segmentation took place.

Whilst the simulated distributions do not obtain an exact match with experimental results for these shapes, there are nonetheless indicators that the modelling assumptions have potential to explain the segmentation. As can be seen in (Fig 5 IB and IIB), the simulated modes shift away from the wFPM for this set of shapes in the direction of the sFPM. On the other hand, in (Fig 5 IIIB), the mode is closer to the wFPM than the sFPM, as expected from the experimental results. Given that these are “raw” results, obtained with no parameter tuning, the assumptions used in this modelling work could suffice to explain the apparent image segmentation. Still, we do not exclude that further dynamics are needed to fully capture these experimental results (see Discussion).

Interestingly, the model predicts a mode at the correct location in (Fig 5 IC, IIC), without having to resort to edge (or feature) matching, which was the explanation given in the original paper of Lent et al (2013a). Over all sets of examples, the pooled distributions obtain an average distance between modes of 11.6° ± 11.5° (mean ± std). When studying the performances over all shapes and parameters separately we obtain a distance between modes of 10.1° ± 9.3° (mean ± std). The average performance against parameter values are shown in (S2 Fig Panel C). Whilst the simulated distributions for composite shapes have lower performance and higher variance than those for previous sets of shapes, the results of the computational model are not inconsistent with the experimental data, and show signs that the model can capture several aspects of the experimental findings.

## Discussion

Visual memory based navigation in insects has been extensively studied at a behavioural level [2–5, 40], in models [9, 12, 14, 15], and in terms of its underlying neural substrates [16–18, 41, 42]. Whilst it is clear that ants collect and use visual memories to navigate their environment, it is still largely unknown what aspects of a visual scene are memorized and how the memories are used to affect steering decisions [20, 21, 33, 43]. Experiments have shown that the wood ant *Formica rufa* can learn a heading direction using the proportion of a large shape that lies left of its visual centre in training [22], a proportion referred to as fractional position of mass (or FPM for short). This suggests that ants can memorize higher-order features extracted over the whole extent of a visual scene. In this work, we used a simple model leveraging known neuroanatomy and connectivity in ants [35–37] to explore whether navigation to the FPM could be a resulting factor of the bilateral structure of the visual and memory circuits. In so doing, we assumed a retinotopic image representation, and showed how the FPM of a visual scene can be implicitly encoded in the memory dynamics of the bilateral network. This work therefore demonstrates that some higher order visual processing and visual cognition could be emergent properties of the overall neural architecture rather than outputs of discrete visual processing modules.

### What visual processing is underpinning visual navigation in ants?

It is still unknown what visual information is used by ants in visual memory based navigation. At a neural level, visual information reaches the MBs via the anterior superior optical tract (ASOT) which relays information from each visual field to both MBs. Neuroscientific studies [35–37] reveal extensive projections from the Medulla (ME) of the Optic Lobes (OL) to the MBs, and a smaller amount of projections from the Lobula (LO). The ME and LO are known to specialize respectively in small and wide field information and can convey chromatic, temporal, and motion features in bees [44]. A range of experimental studies [22, 24–27, 45] highlight that insects use higher-order features extracted over the whole visual scene, such as the center of mass (CoM) or fractional position of mass (FPM) of large shapes. However, many of these results can be explained by the present study, or by other models showing higher-order parameters emerging from a population of simple visual filters [46].

The success of this modelling approach relied upon the assumption that visual processing preserves sufficient retinotopy so that ratios are roughly conserved from retinal to vPN layers (see theoretical section of results). To clarify, we do not need the vPNs to have an exact topographic representation of the visual scene, but to preserve a statistical representations of the retinal information. In this sense, this approach may be stable against the processing by different kinds of filters which may occur within the ME and LO of the Optic Lobes, which we can assume to preserve a kind of semi-retinotopic representation. On the other extreme of the representation spectrum are approaches that are composed solely of features in the global spatial frequency domain, such as rotationally invariant features based upon Zernike moments [28, 29]. We do not expect our modelling approach to work if we apply such processing because this type of representation would not preserve the statistics of the retinotopic representation.

In modelling, we further assumed that the visual Projection Neurons (vPNs) respond to the black parts of the visual scene which defines the shape. This is consistent with experimental and modelling papers that suggest ant photoreceptor tuning might pre-process visual scenes into objects that contrast against sky [47]. Graham and Cheng (2009) show the importance of the panoramic skyline, of objects appearing as dark regions in a scene contrasted against the bright sky. Similarly robot studies have shown the importance of sky segmentation (based on UV light) for robust navigation across different lighting conditions [48, 49]. Together, these studies suggest that the early stages of the ant’s visual system may be optimized for separating the sky from ground, leading to a more reliable image representation (e.g., invariant to weather patterns).

### Is segmentation the result of a selective attention mechanism?

The modelling assumptions of the present study partially helped in explaining the experimentally obtained heading directions for composite shapes, which hinted at a visual segmentation mechanism. However, it is possible that further dynamics are needed to fully explain these results. One such candidate is that ant vision is endowed with an attention-like process [50]. Whilst attention is a concept more commonly associated with primates [51, 52], attention-like processes have also been demonstrated in several insect species [50, 53]. Visual selective attention plays a fundamental role in tracking, whereby a salient target can suppress the responses of other targets, as shown in e.g., praying mantises [54], fruit flies [55] and dragonflies [56]. However, it is not known whether such attentional processes would also be used in visual navigation.

If the ant visual system selectively favours information coming from a structurally cohesive and central part of the visual scene, then the visual memories could be limited to one of two triangles explaining the effective segmentation of the visual scene. As mentioned above, this upward weighting of one triangle over the other could be the result of a selective attention mechanism. However, we can also imagine obtaining similar results from an architectural mechanism, for instance by having higher density of projections attributed to the central parts of the visual fields. Another part of MB dynamics that was not considered in our modelling work is the existence of recurrent connections at the Kenyon Cell layer [57]. These connections have the potential to complexify the memory formation process, for instance they have been shown to allow sequence learning [58] and point attractor dynamics [59]. It is possible that visual information undergoes a transformation due to the KC to KC recurrent connections which dampens peripheral information, providing another speculative explanation for the effective segmentation of the visual scene.

### How do the MBONs drive downstream steering behaviours?

In this modelling work, we explored whether a signal for the FPM-match direction exists within the dynamics of bilateral memories. In so doing we did not make assumptions as to how that signal may be used downstream to drive behaviour, and therefore the model could be used in a range of visuo-motor control strategies. However, considering the specific behaviour that was used in the original paper to measure goal directions can lead to interesting speculations into it’s neural mechanism. The goal directions in Lent et al (2013a) are measured using the end-points of saccade-like-body turns (SLTs), which are periods of high rotational velocity shown in [39] to be visually guided turns correcting the heading direction towards a goal. The fact that the angular speed of saccades is proportional to their angular error at the beginning of the turn [39] is a strong indicator that ants have an internal estimate of turn size prior to initiating saccades. This finding shows striking similarities with the recently discovered function of PFL2 cells in *Drosophila melanogaster*, which have been demonstrated to adaptively tune the speed of turns based on the angular difference between a goal direction and the current heading direction maintained within the Central Complex [60]. The integration of contextual (body-centric) information with a world-centric reference frame (and vice versa) has been shown to rely upon the Fanshaped Body (FB) of the Central Complex (CX) in numerous studies [60–64]. In light of these facts, we postulate that in ants, Mushroom Body outputs (MBONs) are used to set a goal direction within the FB of the CX. Because of the conclusions of our present study, we speculate that goal directions are set when the novelty is (comparatively) low and also balanced across left and right hemispheres. Saccades would then constitute turns that correct the current heading direction towards this set goal direction.

The fact that SLTs occur at a specific phase [65] within the intrinsic oscillatory cycle of the ants [66], further suggests a recurrent signal from the Lateral Accessory Lobes (LALs) is implicated in initiating the turns. This fits well with the dense connectivity patterns between the FB and LALs forming functional loops [67, 68]. We further note that since the turns occur at the outer edge of the oscillations [65], the difference between the current heading and the goal direction constitutes a prediction error (this difference should be zero if ant is on track). Therefore, it seems that SLTs constitute a correction mechanism scaffolded on top of other navigational strategies.

## Conclusion

The major modelling insight and prediction of this study, is that a bilateral brain architecture allows ants to balance the quality of their memories in left and right hemispheres when navigating a simple visual scene. As well as allowing ants to implicitly use higher-order visual properties for navigation, it is possible that having two separate assessments of a view contributes to the robustness of the entailed behaviors [32]. Still, we do not known the full extent of how the bilateral brain architecture contributes to navigational functions.

## Methods

### Image Geometry

The experimental setting of Lent et al (2013a) consisted of an arena of radius *R* = 1.5*m* within which ants were trained to head to a feeder *r* = 0.6*m* away from the centre (Fig 1A). The stimulus angles of the shapes used (as measured from the center of the arena) are given in the original paper, and inferred where exact values are not given. Images are exported with vertical and horizontal axes corresponding respectively to polar and azimuthal angles. We represent the black color with pixel value 1 and white with pixel value 0, allowing for grey scale values in between.

The equations for how the experienced shape transforms as a hypothetical ant moves within a circular arena were determined using trigonometry. Let *Z* be the point at the base of the arena wall representing azimuthal angle 0°. Let *θ* and *ϕ* be the azimuthal and polar stimulus angles of a point *P* on the arena wall as measured from the center *O* of the arena. If *L* is a new location within the arena and *P*_*xy*_ = (*R* cos(*θ*), *R* sin(*θ*)) is the location of point *P* projected onto the base of the arena, then the angles transform as follows:

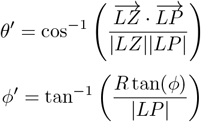

By transforming all points constituting the outline of a shape, we export (90 by 360) black and white images representing the visual pattern as experienced at required locations in the arena.

### Image Processing

From an exported image (panoramic), we extract two images corresponding to the left and right visual fields, according to an overlap parameter (Fig 1B). We do so by cropping the azimuthal angles to the range 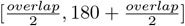 for the left visual field, and range 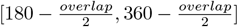 for the right visual field. The two images are then stacked horizontally creating one feature-image with two implicit parts. To better capture the resolution of ant vision, we reduce the image resolution by cropping the top 10 layers of pixels, and downsampling the resulting image by a factor of 4. The downsampling is done by taking the mean of each 4 by 4 block in the (80 by 360) cropped image, resulting in a (20 by 90) image with 4° per pixel resolution.

The training paths are created assuming an ant moves directly from the centre of the arena towards the feeder location, facing straight ahead. We export *N*_*train*_ = 30 images for each training pattern taken in equal steps from distance *r* = 0*m* to *r* = 0.6*m*. Given that we cannot assume in advance a movement direction for simulated trajectories, we only use *N*_*test*_ = 1 image taken from the centre of the arena for the test shapes.

### Network Architecture

We model the Mushroom Bodies with an artificial neural network (ANN), represented pictorially in (Fig 1B). The model’s implementation details are most similar to those used by Le Moël and Wystrach (2024). The network is composed of three layers with four types of neurons:

- The first layer is the input image corresponding to the left and right visual fields stacked horizontally, which is a matrix of shape (20, 90). The Visual Projection neurons (vPNs) fire with strenght given by the grey scale value of the corresponding pixel ranging between 0 (white) and 1 (black).
- The second layer is composed of two Mushroom Body (MBs) compartments, each having *N*_*KC*_ = 25, 000 Kenyon Cell (KC) neurons. The input strength of a KC is computed as a weighted sum over connected pixel values input(*KC*) = Σ_*x*_ *w*_*xkc*_ × *x*. Where *w*_*xkc*_ denotes the weight of the connection from pixel *x* to the given KC (as described in next section).
- Each MB compartment is also fully connected to an Anterior Paired Lateral neuron (APL), which induces a normalisation on the overall activity of the MB [69]. For a given pattern of firing Kenyon Cells, the APL will select the proportion *p* of cells with the strongest activity and allow them to fire (output is set to 1), all other cells do not fire (output is set to 0). In essence, this binarises Kenyon Cell activity and ensures a sparse representation.
- The third layer is composed of two Mushroom Body Output Neurons (MBONs), which fire as a weighted sum of Kenyon Cell outputs: output(*MBON*) = Σ_*KC*_ *w*_*KC*_ × output(*KC*). The connections and weights between Kenyon Cells and MBONs are described in the next section.

### Network Connections

The layers are connected and weights are set as follows:

- A sparse pattern of connections is created between vPNs and KCs so that KCs have on average *K* connections with vPNs. The connections are generated by picking 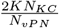 Kenyon Cells for each vPN according to the following bias: probability 1 − *crossover* of connecting to the ipsilateral MB, and probability *crossover* of connecting with the contralateral MB.
- We consider two modes of weight initialization. In the first case (constant weights), all vPN to KC connection weights are initialised to *w*_*x*_ = 1*/K*. In the second case (random weights), each weight is assigned a random and uniform value in the range 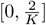. chosen to have a maxima that weights have a mean value of 1*/K* in both cases.
- Each MB is fully connected to the MBON ipsilateral to it. Connection weights are initialised to 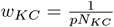, chosen to have a maximal expected novelty value of 1. As with the previous case, we have a random mode of weight initialization where each output weight is assigned a random and uniform value in the range 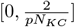 These weights are subsequently updated during training, as explained in the next section.

### Network Training

The training acts solely on the connections from Kenyon Cells to MBONs, and works by depreciating the weights from KCs that fire for the train image(s). This type of update is often referred to as “Anti-Hebbian” despite no explicit referral to the postsynaptic neuron’s firing. In Hebbian learning, neurons that fire in close temporal proximity would have their synaptic weights strengthened (in Anti-Hebbian, synaptic weights are weakened). In our model, the action of Dopaminergic neurons (DANs) and the post synaptic neuron’s firing (MBON) at train time are not explicitly modelled, making for a simpler learning rule which considers only Kenyon Cells. Each time a Kenyon Cell fires in the training phase, its output weight *w*_*KC*_ is multiplied by a factor of *α <* 1.

The weight update for the whole training can be condensed into a single formula. In effect, each weight is multiplied by a factor of 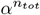 where *n*_*tot*_ is the total number of times the KC fired in training. In equations:

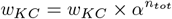

We use *α* = 0.95 as the default value and study the effect of the parameter in the supplemental material.

### Network Testing

To test a network with a single image, we forward-pass the image through the trained network and take the output of the MBONs (Fig 1B) as measures of (left and right) novelties. To produce rotational signals, we forward-pass the image associated with each of the 360 possible facing directions. With two MB outputs, there are four signals that we consider: *left novelty, right novelty, novelty sum, novelty difference* (S1 Fig). The novelty sum and difference are used to produce distributions of goal directions, as described in the next section. Rotational signals are displayed as the mean of *n*_*models*_ = 50 model initializations, with a shaded area spanning one standard deviation either side of the mean (S1 Fig, S3 Fig).

### Generating Heading Distributions

From the rotational novelty difference and rotational novelty sum signals (S1 FigC-D), we extract distributions that model the distribution of ant saccade-like-turn (SLTs) end points from Lent et al (2013a). To produce a distribution from the outputs of our computational model, we take a Monte Carlo approach. The method assumes that both proximity to the minimum of the novelty sum and proximity to a zero-crossing of the novelty difference are important for determining goal directions. We generate tentative directions uniformly at random in range [−180°, 180°] and accept with probability:

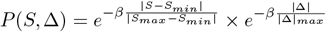

Where *S* indicates the novelty sum for that given direction, and Δ is the novelty difference. *S*_*max*_, *S*_*min*_ are the maximum and minimum values of the novelty sum, |Δ| _*max*_ is the maximum absolute novelty difference (see S1 FigC-D). The value of the exponential decay *β* tunes the variance of the simulated distributions, and we use *β* = 4 to qualitatively match the variance of the distributions in Lent et al (2013a). Histograms are produced by binning the data in range [−70, 200] with a bin width of 5°, to match the presentation of results in Lent et al (2013a).

### Pooling Heading Distributions

Given a desire to be sure that our results are derived mainly from the architecture of the model, rather than esoteric parameter choices, we generate the aggregate behaviour of the model by pooling results over a range of parameter settings. Parameters *α* = 0.95, *K* = 8, *p* = 0.05 are kept fixed at their default values because they have little effect on model outputs as far as zero-crossings are concerned (S3 Fig), and we systematically study all 30 cross pairs of *crossover* and *overlap* in the ranges [0, 0.1, 0.2, 0.3, 0.4] and [0°, 8°, 16°, 24°, 32°, 40°] respectively. The range of overlap values was chosen based on the experimentally observed anatomy of *Cataglyphis sp*. in [70].

For each of the 5 *crossover* parameter values, 50 models are initialized. Each of the 50 models is trained in turn with each of the 6 visual *overlap* parameter values. The resulting 50 × 30 model and training combinations are independently tested to produce rotational novelty difference and rotational novelty sum signals, which are used to simulate *n*_*saccade*_ = 100 SLT end points each (as described in previous section). We therefore pool together *n*_*saccade*_ = 100 saccade end points for *n*_*repeat*_ = 50 model initializations and *n*_*params*_ = 30 cross pairs of parameter settings, resulting in a total of *n*_*montecarlo*_ = 150, 000 simulated saccades which we use to infer a distribution.

### Distribution Modes

To infer mode locations for the simulated distributions (whilst avoiding small sample artefacts), we use a continuous estimate of our empirical distributions. We produce such a smoothed approximation using Kernel Density Estimation (KDE) with a gaussian kernel, and a bandwidth of *h* = 5° to match the angular acuity of our data. We extract the KDE estimate 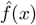 in steps of 1° and define a point x to be a mode if it is a local optima of the KDE function. To avoid duplicate optimums, we ensure a local optimum is preceded by an increase for 5 degrees and succeeded by a decrease for 5 degrees.

### Distribution Performance

For a simulated distribution, we extract a measure of performance (which we refer to as “distance between modes”) using the minimum distance between its modes and the experimentally observed mode, inferred from Lent et al (2013a). In equations

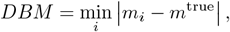

where *m*_*i*_ are the modes extracted from the simulated distribution and *m*^*true*^ is the true mode from the original experiments. We always repeat performance estimations *n*_*iter*_ = 10 times and average results to mitigate for variance. When we compute a performance over *N* experiments, we use the Mean Absolute Error (MAE) of the predictions for the individual experiments:

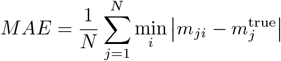

The performances are computed independently for the pooled distributions and for all combinations of *crossover* and *overlap* parameter values (see S2 Fig).

### Model Parameters

The model parameter are listed below along with their function:

- *α*: the learning rate of the network. Controls how fast weights decrease in training.
- *K*: expected number of connections of Kenyon Cells. Controls the sparsity of the network.
- *p*: the proportion of KCs that are allowed to fire. Controls the normalisation of MB activity.
- crossover: the probability that a connection crosses over from one visual field to the contralateral MB. Controls the ipsilateral bias of the network initialization.
- overlap: the amount of angular overlap of left and right visual fields. Changes the azimuthal cutoffs of left and right fields of view.

When a parameter is not explicitly studied, its value reverts to the default given in Table 1. The effect of parameters on model outputs is explored in the supplemental material (S3 Fig).

**Table 1.** Default parameter values and ranges studied.

| Parameter | Default Value | Range Studied |
| --- | --- | --- |
| $\alpha$ | 0.95 | 0.9, 0.95, 0.99 |
| $K$ | 8 | 4, 8, 16, 32 |
| $p$ | 0.05 | 0.025, 0.05, 0.1, 0.2 |
| *crossover* | 0.2 | 0, 0.1, 0.2, 0.3, 0.4, 0.5 |
| *overlap* | 0° | 0°, 8°, 16°, 24°, 32°, 40° |

## Supporting information

**S1 Fig. Signals of bilateral memory network**. The training image used for this example is a rectangle of width 160° and height 38° with feeder inset 30° from the left side of shape (Fig 2A). The test shape is a rectangle of width 80° and height 38° (Fig 2B). (A) The left novelty is plotted as a function of angular orientation, with 0° being the direction to the left edge of the train and test shapes. (B) The right novelty is plotted as a function of angular orientation (C) The sum of left and right novelties is plotted. *S*_*min*_ denotes the minimum value of the signal. *S*_*max*_ denotes the maximum value of the signal. *S* denotes the value of the signal at an angular orientation of 75°. (D) The difference of left and right novelties is plotted against angular orientation. Δ_*max*_ denotes the maximum novelty difference in absolute value. Δ denotes the novelty difference at an angular orientation of 75°.

**S2 Fig. Model performance heatmaps**. Average performance of the model, denoted as DBM (“distance between modes”) heatmaps for all combinations of crossover [0, 0.1, 0.2, 0.3, 0.4] and overlap [0°, 8°, 16°, 24°, 32°, 40°] parameter values shown for (A) First set of experiments. (B) Second set of experiments. (C) Third set of experiments. (D) Overall.

**S3 Fig. Parameter scans for simple train to test comparisons**. The training image used for all results in this figure is a rectangle of width 160° and height 38°. Column (A) consists of testing on a 120° by 38° rectangle. Column (B) consists of testing on a 80° by 38° rectangle. Column (C) consists of testing on a 40° by 38° rectangle. Row (i) consists of rotational novelty difference plots for different values of parameter *α* in the range [0.9, 0.95, 0.99]. Row (ii) studies parameter *K* in range [4, 8, 16, 32]. Row (iii) studies parameter *p* in range [0.025, 0.05, 0.1, 0.2]. Row (iv) studies parameter *crossover* in range [0, 0.1, 0.2, 0.3, 0.4, 0.5]. Row (v) studies parameter *overlap* in range [0°, 8°, 16°, 24°, 32°, 40°]. Row (vi) studies the model with randomly initialized connection weights. In the first case we initialize vPN to KC weights uniform in [0, 2*/K*]. In the second case KC to MBON weights uniform in [0, 2*/pN*_*KC*_]. In the third case, both are initialized randomly.

